# Building a Reference Transcriptome for the Hexaploid Hard Fescue Turfgrass (*Festuca brevipila*) Using a Combination of PacBio Iso-Seq and Illumina Sequencing

**DOI:** 10.1101/2020.02.26.966952

**Authors:** Yinjie Qiu, Ya Yang, Cory D. Hirsch, Eric Watkins

## Abstract

**Background:** Hard fescue (*Festuca brevipila* Tracey, 2n=6x=42) is a cool season turfgrass with a fine leaf texture that performs well under low-input management. Breeding and genetics studies of *F. brevipila* have been limited due to the complexity of its hexaploid genome. To advance our knowledge of *F. brevipila* genomics, we used PacBio isoform sequencing to develop a reference transcriptome for this species.

**Results:** Here, we report the *F. brevipila* reference transcriptome generated from root, crown, leaf, and inflorescence tissues. We obtained 59,510 full-length transcripts, of which 38,556 were non-redundant full-length transcripts. The longest and shortest transcripts were 11,487 and 58 bp, respectively. Distribution of synonymous distances among paralogs within *F. brevipila* suggested highly similar subgenomes that are difficult to distinguish from sequencing errors. To evaluate the phylogenetic relationships among *F. brevipila* and close relatives, we sequenced three additional transcriptomes using closely related species on an Illumina platform. The results of our phylotranscriptomic analysis supported the close relationships among *F. brevipila* (6x), *Festuca ovina* (4x), *Festuca ovina* subsp. *ovina* (2x), and *Festuca valesiaca* (2x), with high levels of discordance among gene trees.

**Conclusions:** Overall, the *F. brevipila* PacBio Isoseq reference transcriptome provided the foundation for transcriptome studies and allowed breeders a resource for gene discovery in this important turfgrass species.

## Background

*Festuca* (*Festuca* L., Poaceae) is a species-rich genus that includes more than 450 species (Clayton and Renvoize, 1986). Several species in this genus have been used as ground cover turfgrass: (1) broader-leaf fescues that includes tall fescue (*Festuca arundinacea*) and meadow fescue (*Festuca pratensis*) and (2) fine-leaf fescues (Wilkinson and Stace, 1991). Five fine-leaved fescue taxa are of particular interest to turfgrass breeders because of their performance in shade, drought tolerance, and adaptation to low fertility and acidic soils (Carroll, 1943; Hanson and Juska, 1969; Reiter et al., 2017). These five fine fescue taxa include hard fescue (*Festuca brevipila* Tracey, 2n=6x=42), sheep fescue (*Festuca ovina*, 2n=4x=28), both of which belong to the *Festuca ovina* complex; and the *Festuca rubra* complex that includes strong creeping red fescue (*Festuca rubra* subsp. *rubra* 2n=8x=56), slender creeping red fescue (*Festuca rubra* subsp. *littoralis*, 2n=6x=42), and Chewings fescue (*Festuca rubra* subsp. *fallax*, 2n=6x=42; Huff and Palazzo, 1998). A previous phylogenetic study using plastid genomes resolved the relationships among these five taxa (Qiu et al., 2019). Of the five fescue taxa, *F. brevipila* is a particularly attractive target for breeding and genetics improvement due to its excellent performance in low-input environments (Ruemmele et al., 1995).

Modern breeding programs in model crop species rely heavily on the use of genomic and molecular biology tools, such as marker-assisted breeding and CRISPR-Cas 9 based genome editing to improve consumer and field production traits (Varshney et al., 2005; Gupta et al., 2010; Shan et al., 2013; Phing Lau et al., 2016). To apply these tools in turfgrass breeding, there is a need to build genomic and transcriptomic resources. However, adapting molecular breeding techniques and genomic studies into *F. brevipila* has been challenging because of a lack of genetic information. As of September 2019, there were 174 nucleotide sequences for *F. brevipila* on National Center for Biotechnology Information (NCBI, www.ncbi.nlm.nih.gov), while perennial ryegrass (*Lolium perenne*), the turfgrass species for which the most genomics work has been done, has a reference genome available, 45,149 Sequence Read Archive (SRA) accessions, and more than 600,000 nucleotide sequences on GenBank.

RNA sequencing is a powerful tool to generate genomic data for marker development and enables researchers to study the dynamics of gene expression of plants under stress. As the cost of Illumina sequencing continues to drop, more and more researchers are using this approach for gene discovery and gene identification. Without a reference genome, researchers use *de novo* assembled short reads to generate a reference to study gene expression (Grabherr et al., 2011). However, this approach is error-prone in polyploids with highly similar subgenomes (Chen et al., 2019), making it difficult to obtain full-length transcripts or identify alternative splicing (Alkan et al., 2011; Ozsolak and Milos, 2011).

PacBio sequencing, also known as long-read sequencing, has average read length over 10 kb (Rhoads and Au, 2015) and is particularly suitable for generating reference transcriptomes in polyploid species. With the capability to obtain full-length RNA sequences (Isoseq), PacBio Isoseq has been used for the discovery of new genes, isoforms, and alternative splicing events; this technology is improving genome annotation in crop species such as maize (*Zea mays*), rice (*Oryza sativa*), sugarcane (*Saccharum officinarum*), and sorghum (*Sorghum bicolor*) (Wang et al., 2016; Hoang et al., 2017; McCormick et al., 2018; Zhang et al., 2019). In turfgrass and ground cover related genetics research, PacBio Isoseq has been used to study the transcriptome and phylogeny of bermudagrass (*Cynodon dactylon*) and red clover (*Trifolium pratense*) (Chao et al., 2018; Zhang et al., 2018). Due to the highly outcrossing and polyploid nature of *F. brevipila*, PacBio isoform sequencing is ideal for generating a reference transcriptome for plant breeding and genetics research.

For plant breeding and germplasm improvement, USDA PI accessions have proven to be a valuable resource (Pastor-Corrales, 2003; Chang and Hartman, 2017). Our previous work found that out of 127 USDA accessions of the closely related *Festuca ovina*, 17.3% are hexaploid, 33.1% are diploid, and 26.8% are tetraploid (Qiu et al., 2020a). For this reason, it is important to understand the genomic content of our focal species so that breeders can potentially create synthetic hexaploids using taxa with lower ploidy levels for traits introgression. Previous analyses on GC content, average chromosome size, and genome size suggest the presence of allo/auto-polyploids across *Festuca* (Šmarda et al., 2008). However, there is a lack of nuclear sequence-based evidence to resolve the phylogenetic relationships of *F. brevipila*. Previous research using the amplified fragment length polymorphism (AFLP) markers followed by neighbor-joining analysis suggested that *F. valesiaca* (2n=2x=14) is closely related to *F. brevipila* (Ma et al., 2014). By sequencing the chloroplast genomes, our previous work suggested *F. ovina* subsp. *ovina* (2n=2x=14) and *F. ovina* (2n=4x=28) were both closely related to *F. brevipila* (Qiu et al., 2019). Here we adopt a phylotranscriptomic approach to investigate the phylogenetic relationship among *F. brevipila* and close relatives using nuclear loci.

In this study, we used the PacBio Sequel II platform to conduct isoform sequencing to develop a reference transcriptome for the hexaploid *F. brevipila*. We then evaluated and annotated the reference transcriptome using publicly available protein databases. In addition, to evaluate the nuclear phylogenetic relationships among *F. brevipila* and its close relatives, we used Illumina RNA sequencing on leaf tissue of *F. brevipila, F. ovina* subsp *ovina* (two accessions, one diploid and one tetraploid), and *F. valesiaca*, and carried out phylotranscriptomic analysis together with publicly available genomes and transcriptome datasets.

## Results

### *Festuca brevipila* Transcriptome Assembly

Using PacBio Isoseq, a total of 76.3 GB of sequence was generated from four SMRT cells, 73.91 Gb (48,788,662) of which were subreads. The Isoseq 3 pipeline identified 60,719 putative high-quality full length (FL) transcripts and 487 low-quality FL transcripts. After removing microbial contamination and plastid genome sequences, 59,510 FL transcripts were retained (98.01%) with a N50 of 2,668 bp and a total length of 130,657,908 bp. For Illumina sequencing on *F. brevipila* leaf tissue, a total of 37,100,522 paired-end reads were generated (SRR10995913). The reads were used for PacBio read error correction. Next, we generated the polished *F. brevipila* transcriptome which included 59,510 FL transcripts with a N50 of 2,667 bp and a total length of 130,631,252 bp. After removing redundant sequences by sequence similarity of 99%, we obtained the final reference transcriptome that contains 38,556 (64.78%) unique transcripts with a total length of 82,782,352 bp and a N50 of 2,584 bp. The longest and shortest transcripts were 11,487 and 58 bp, respectively. The GC content of the transcriptome was 51.5%, with the majority of transcripts size above 1,000 bp (83.3%) (**Figure 2)**.

**Figure 1.**
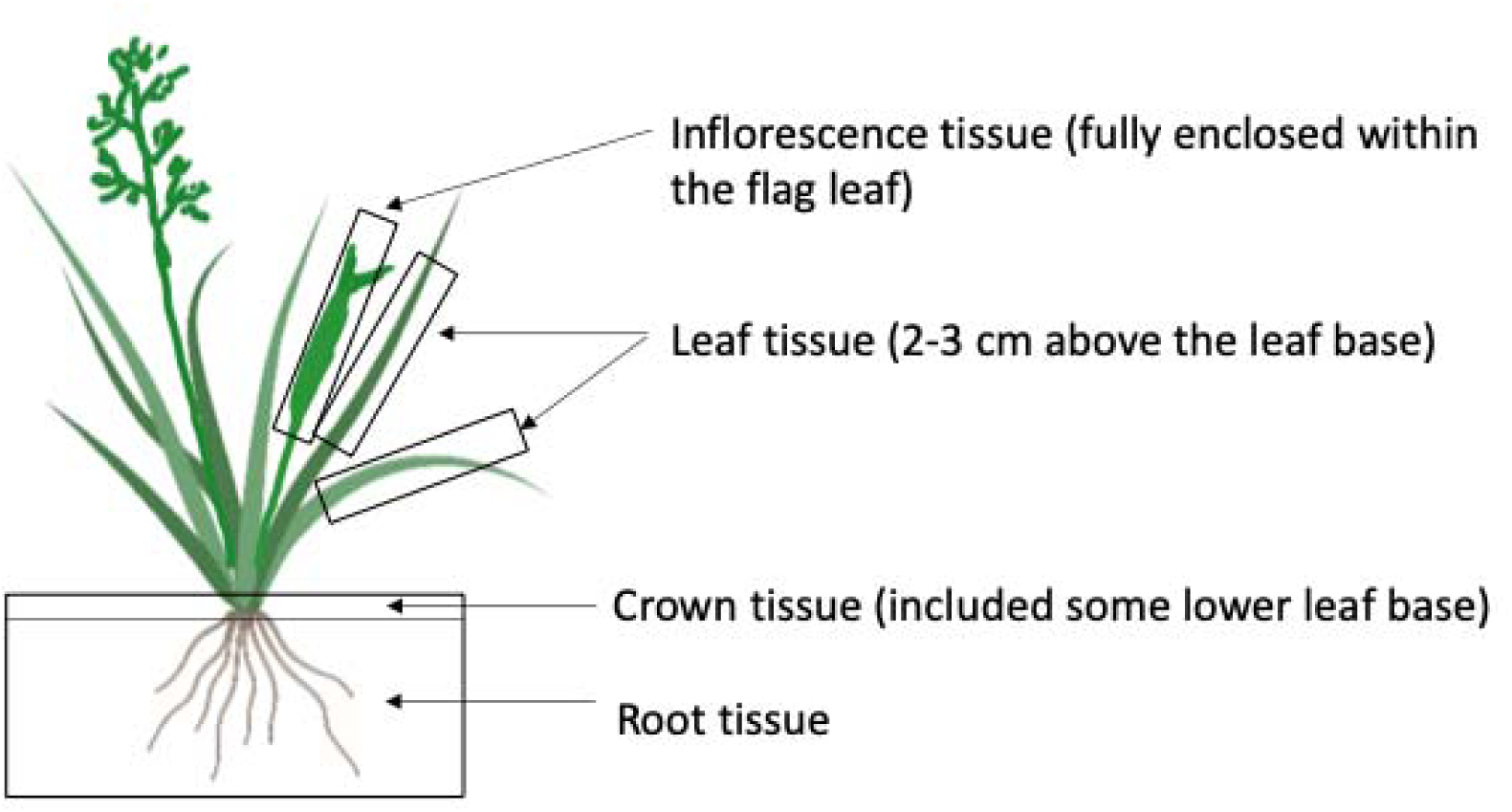
Plant tissues used for PacBio Isoform and RNA sequencing. Because the lower leaf base and crown were not separable, the crown tissue included some lower leaf base. Inflorescence tissue was sampled when it was still fully enclosed within the flag leaf. For the leaf sample, tissue that was 2-3 cm above the leaf base was harvested.

**Figure 2.**
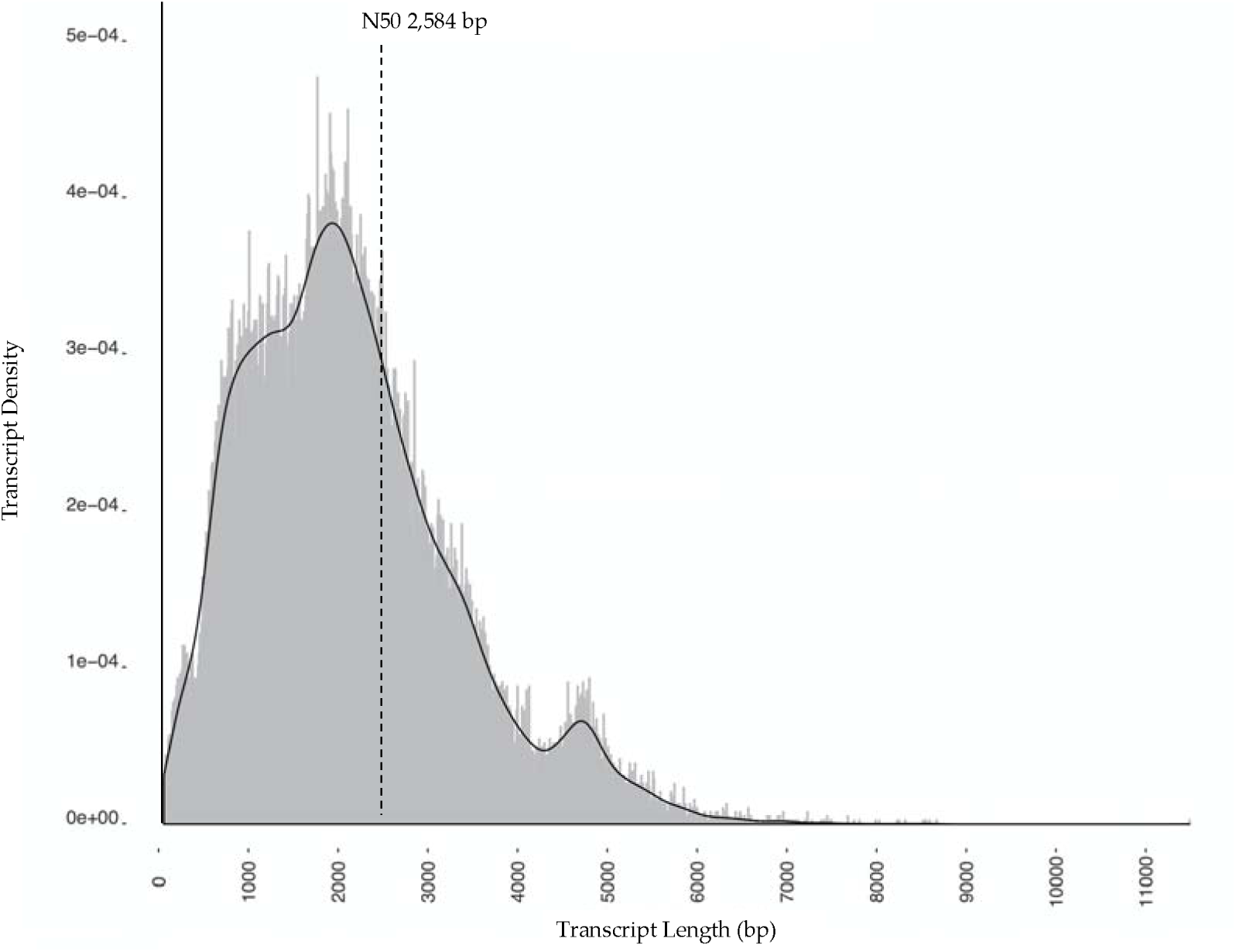
Transcript length distribution after redundant sequence removal.

*De novo* assembly of Illumina reads recovered 325,781 assembled contigs. After contamination and redundancy removal, 302,471 contigs were retained with a N50 of 1,157 bp. The combined assembly of long- and short-read data by 95% sequence similarity included 248,318 transcripts and an N50 of 1,405 bp. Of these, 20,063 transcripts were from the Isoseq dataset (**Table 2**).

**Table 1.**
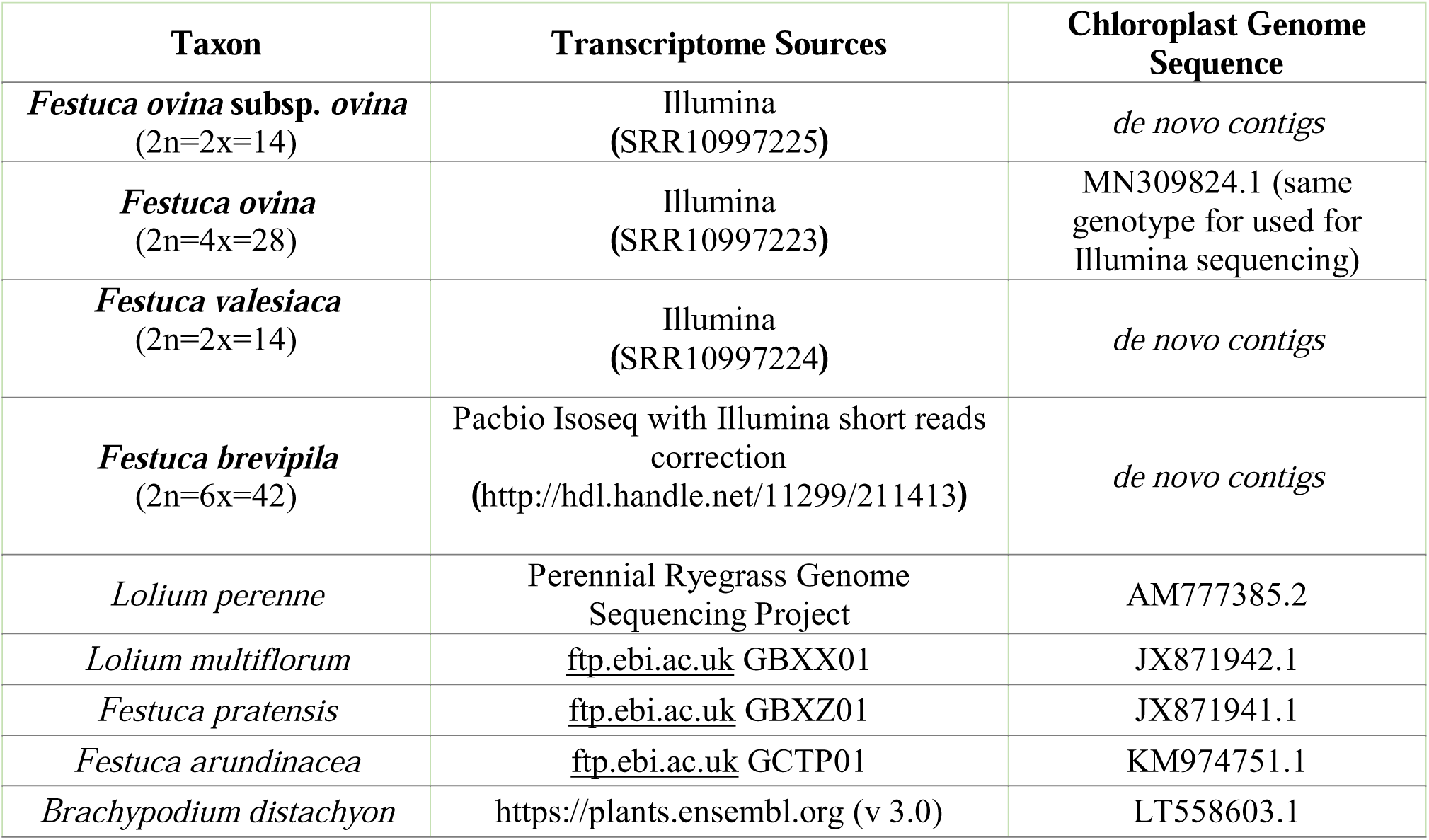
Sources of transcriptome and chloroplast genome sequences used in this study. Sequences newly generated in this study were in bold.

| **Taxon** | **Transcriptome Sources** | **Chloroplast Genome Sequence** |
| --- | --- | --- |
| ***Festuca ovina* subsp. *ovina***<br>(2n=2x=14) | Illumina<br>(SRR10997225) | *de novo* contigs |
| ***Festuca ovina***<br>(2n=4x=28) | Illumina<br>(SRR10997223) | MN309824.1 (same genotype for used for Illumina sequencing) |
| ***Festuca valesiaca***<br>(2n=2x=14) | Illumina<br>(SRR10997224) | *de novo* contigs |
| ***Festuca brevipila***<br>(2n=6x=42) | Pacbio Isoseq with Illumina short reads correction<br>(http://hdl.handle.net/11299/211413) | *de novo* contigs |
| *Lolium perenne* | Perennial Ryegrass Genome Sequencing Project | AM777385.2 |
| *Lolium multiflorum* | ftp.ebi.ac.uk GBXX01 | JX871942.1 |
| *Festuca pratensis* | ftp.ebi.ac.uk GBXZ01 | JX871941.1 |
| *Festuca arundinacea* | ftp.ebi.ac.uk GCTP01 | KM974751.1 |
| *Brachypodium distachyon* | https://plants.ensembl.org (v 3.0) | LT558603.1 |

**Table 2.** Statistics from *F. brevipila* PacBio Isoseq, *de novo* transcriptome, and combined assemblies. No decontamination was carried out for the combined assembly.

| **Assembly** | **Total**<br>**Contigs** | **Contigs after**<br>**Contaminants**<br>**Removal** | **Contigs after**<br>**Redundancy**<br>**Removal** | **N50**<br>**(bp)** |
| --- | --- | --- | --- | --- |
| *de novo* | 325,781 | 322,110 | 302,471 (92.84%) | 1,157 |
| PacBio Isoseq | 60,719 | 59,510 | 38,556 (64.79%) | 2,584 |
| Combined Transcriptome<br>(cd-hit-est) | 341,027 | NA | 248,314 (72.81%) | 1,405 |

When evaluating the completeness of the assemblies, the Illumina *de novo* assembly showed better coverage (94.6%, fragmented BUSCO included) than the PacBio Isoseq transcriptome (71.85%, fragmented BUSCO included). The combined assembly outperformed the individual assemblies for better coverage (96.07%, fragmented BUSCO included) and completeness (89.24%) (**Figure 3**). Around 60% of the fragmented and 35% of the missing BUSCO genes from the Illumina-only assembly were complete in the hybrid assembly (**Figure 4A**). Similarly, 54% of the fragmented and 78% of the missing BUSCO genes from the PacBio assembly were marked as complete in the hybrid assembly (**Figure 4B**).

**Figure 3.**
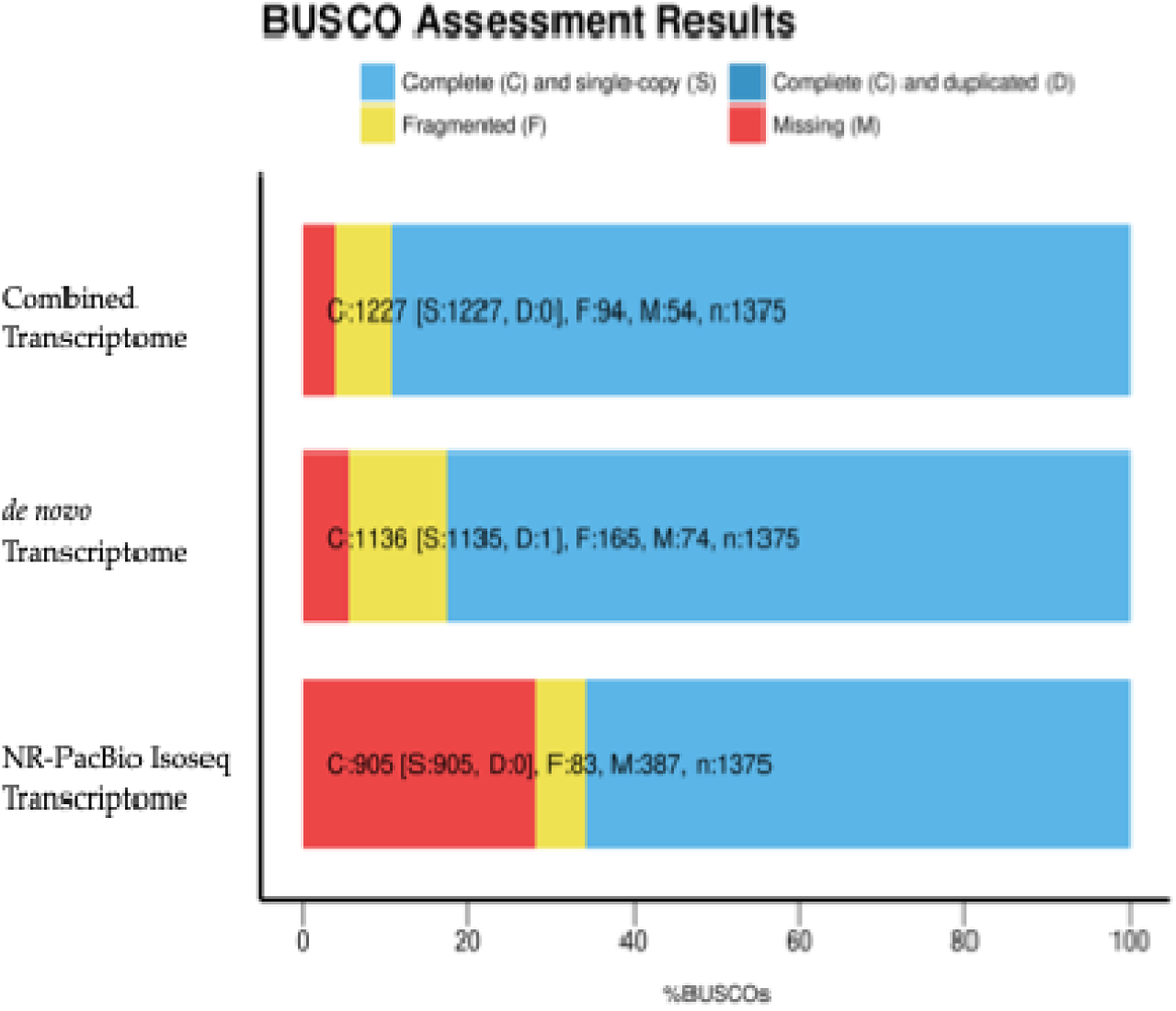
BUSCO assessment of the three assemblies. The combined assembly had the most complete BUSCO genes and the least missing genes.

**Figure 4.**
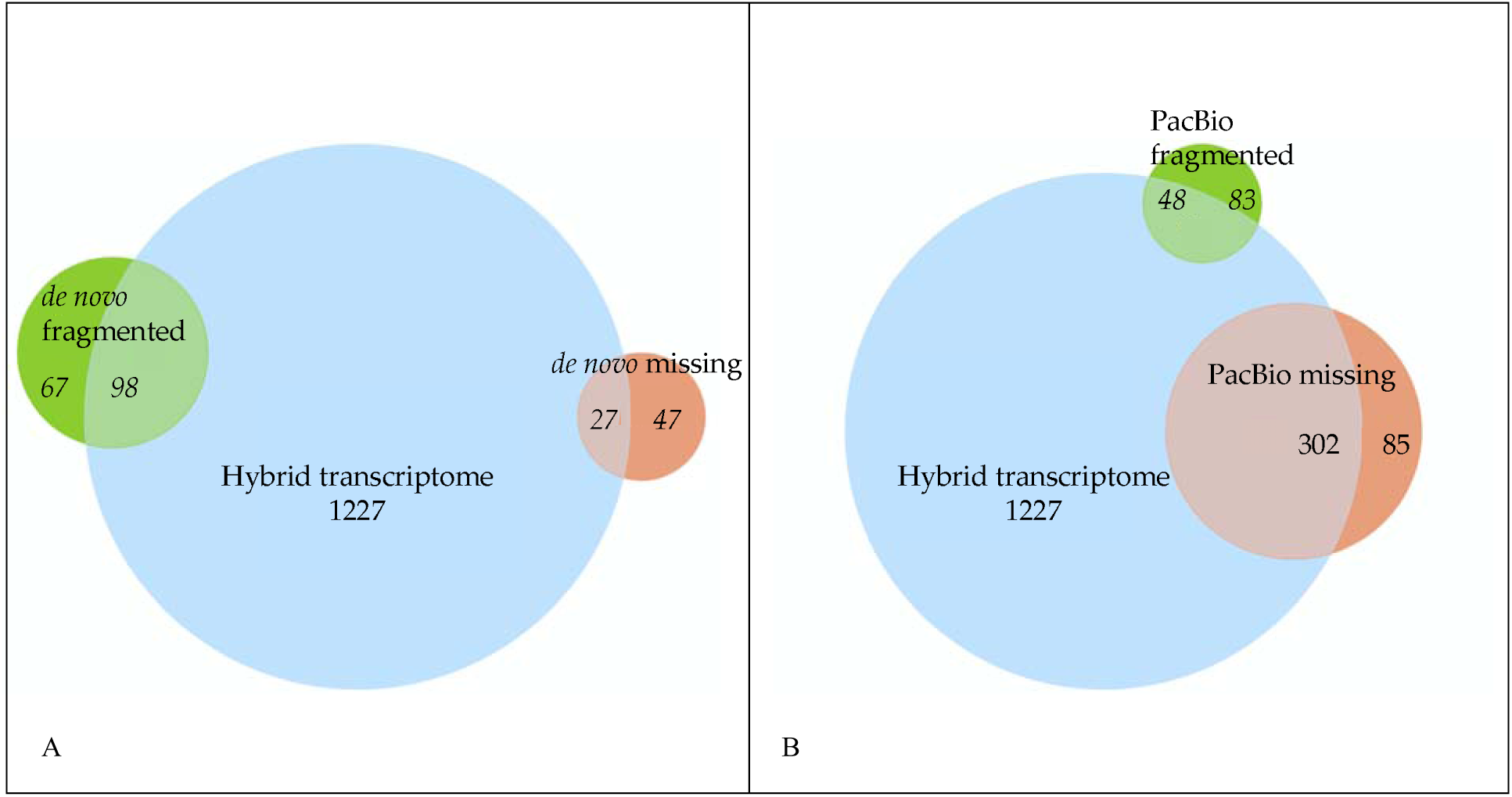
Comparison of BUSCO completeness. Fragmented or missing BUSCO in the Illumina assembly (A) or the PacBio assembly (B) but were complete (1,227) in the hybrid assembly.

When mapping Illumina reads from the *F. brevipila* leaf tissue back to the three assemblies, the percent of unmapped reads for PacBio Isoseq, Illumina, and combined assembly was 34.5%, 38.2%, and 28.5%, respectively.

### Transcript Functional Annotation

We carried out functional annotation of the reference transcriptome using multiple databases including NCBI NR protein, UniRef90, SwisProt, KEGG, and Pfam (GO terms) (**Figure 5**). Around 25% of transcripts had annotation in all five databases and 36,067 (93.54%) of transcripts had annotation from more than one database. The reference transcriptome and annotation have been deposited at https://conservancy.umn.edu/handle/11299/211413 for public access.

**Figure 5.**
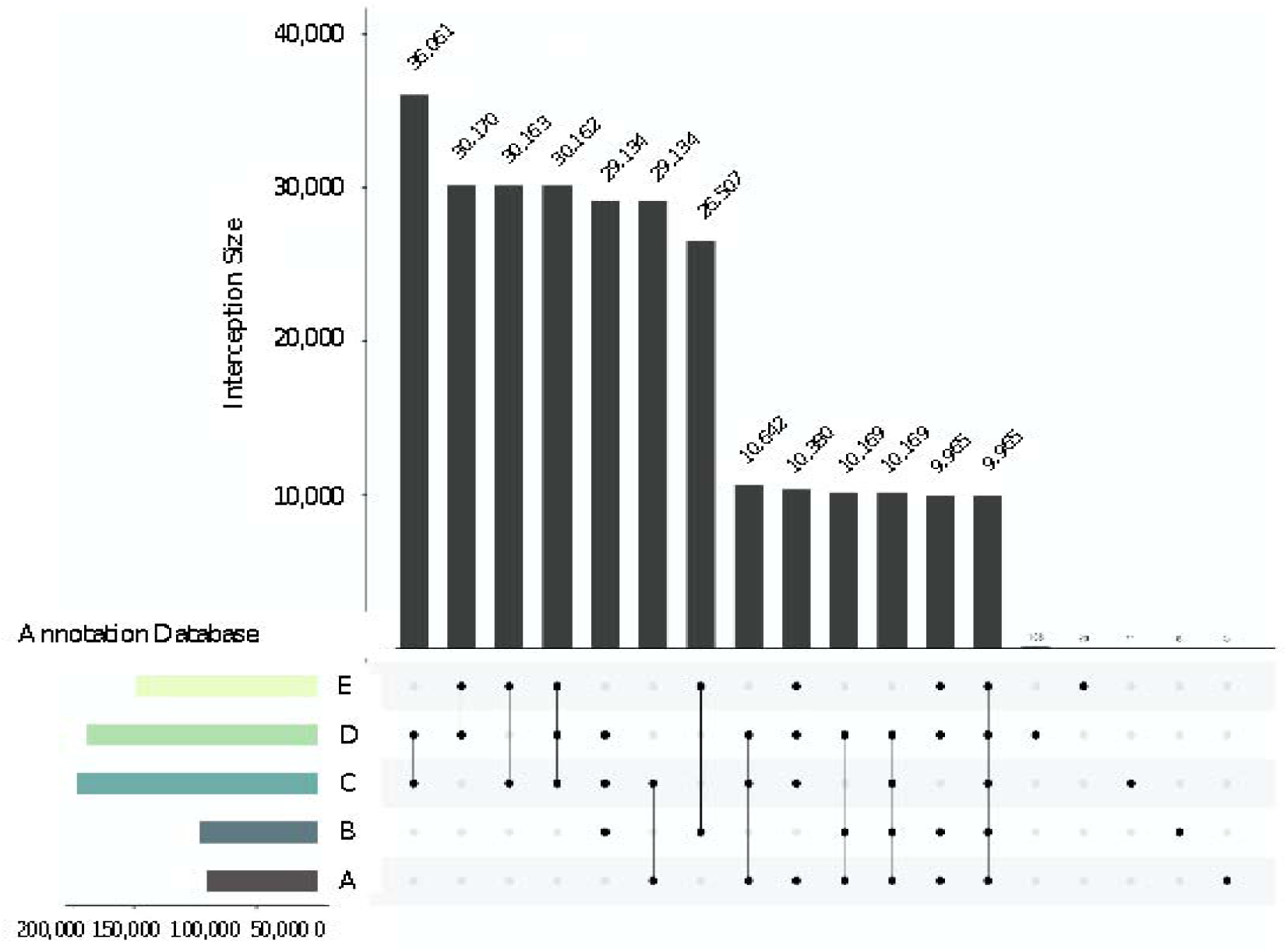
Comparison of *Festuca brevipila* PacBio Isoseq transcriptome annotation using different databases. A: Kyoto Encyclopedia of Genes and Genomes; B: Gene Ontology; C: NCBI NR Protein; D: UniRef90; E: SwisProt.

### Identification of lncRNAs and miRNA from PacBio Sequences

From the Non-Redundant (NR, Isoseq transcriptome after cd-hit-est at 99% sequence identity) PacBio Isoseq reference transcriptome, we identified 7,868 transcripts that were long non-coding RNA. Of them, 145 transcripts were identified as hairpin miRNA, and non-plant hits were removed from further analysis. A total of 39.04% of the miRNA were annotated in *Hordeum vulgare*, 19.18% were annotated in *O. sativa*, and 17.8% were annotated in *F. arundinacea* (**Table S2**). The most copies found was hvu-MIR6179 with 16 copies, followed by far-MIR1119 and osa-MIR2927, both of which had 11 copies (**Figure 6**).

**Figure 6.**
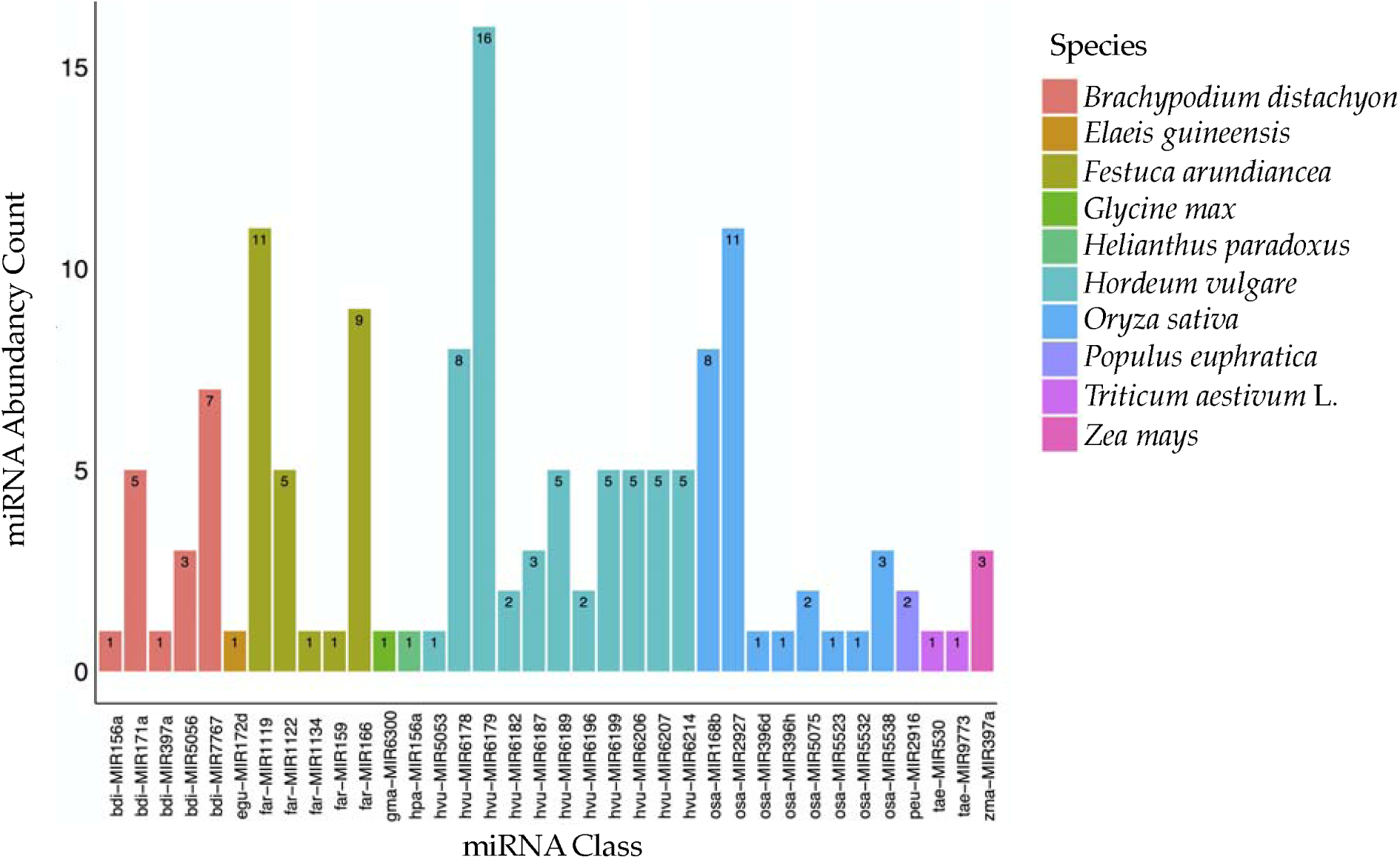
miRNA identified in the PacBio Isoseq reference transcriptome. Most of the homologous miRNAs were identified in barley (*Hordeum vulgare*) and tall fescue (*Festuca arundinacea*). MIR6179, MIR1119, and MIR2927 had the most abundance.

### Phylotranscriptomic Analyses of the *Festuca-Lolium* Complex

A total of 86,445,109 paired end reads were generated from the Illumina sequencing for *F. ovina* subsp. *ovina, F. valesiaca*, and *F. ovina*. The assembly statistics were summarized in **Table 3**.

**Table 3.** Summary statistics of Illumina sequencing and de novo transcriptome assembly. No decontamination was carried out for the three assemblies.

| **Taxon** | **Number of Raw Reads** | **Number of Contigs** | **N50 (bp)** |
| --- | --- | --- | --- |
| *F. ovina* (4x) | 26,640,768 | 242,794 | 1,230 |
| *Festuca ovina* subsp. *ovina* (2x) | 24,719,943 | 179,393 | 1,664 |
| *F. valeciaca* (2x) | 35,084,398 | 157,926 | 1,770 |

OrthoFinder identified 29 single copy nuclear genes across the 9 taxa to reconstruct the nuclear gene trees and three chloroplast genome regions that included 9,680 bp sequence which covered the *rps2-atpA, rpl32*-*ccsA*, and *clpP*-*psbH* regions. These were likely DNA contamination in the RNA sequencing library, because these contigs spanned gene and intergenic regions. The topology between concatenated nuclear and plastome datasets were mostly congruent, except the placement of *F. brevipila* and *F. valesiaca* (**Figure 7**).

**Figure 7.**
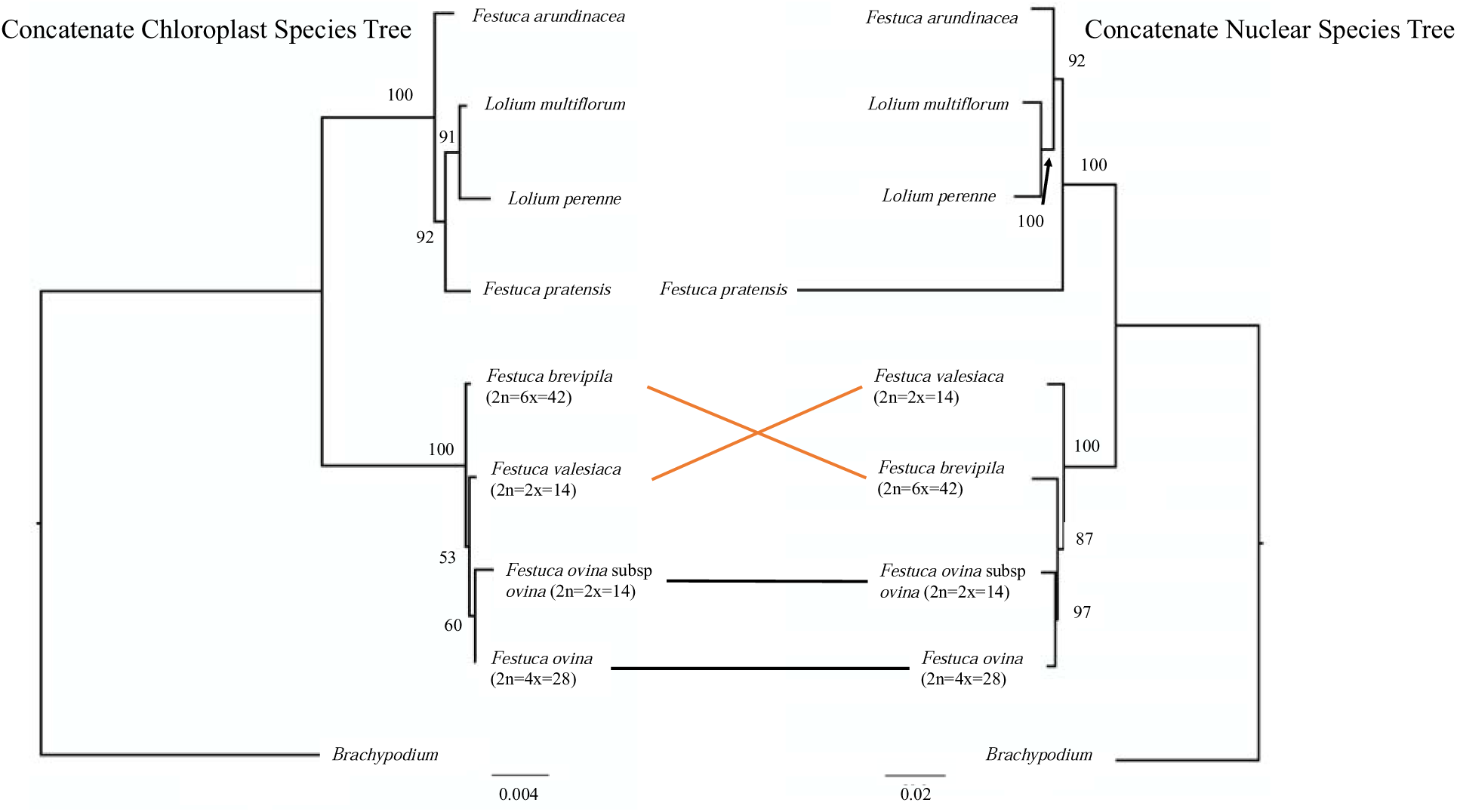
Nuclear and chloroplast species tree of the 9 taxa used in this study. In the *F. ovina* complex, concordance relationship was linked in black while conflicts were linked in orange.

To explore the conflict among gene trees among *F. ovina* (4x), *F. ovina* subsp. *ovina* (4x), *F. brevipila* (6x), and *F. valesiaca* (2x), we assembled a second phylotranscriptomic dataset with these four taxa and two outgroups. OrthoFinder recovered 286 single copy genes from this six-taxon dataset. The resulting species tree topology by ASTRAL was congruent with the concatenated tree recovered from the nine-taxon nuclear dataset (**Figure 8**). However, a significant proportion of well-supported (bootstrap >50) gene trees supported alternative placements of *F. brevipila* (red and green in Fig. 9) being either sister to *F. valesiaca, F. ovina* 2x, or *F. ovina* subsp. *ovina* 4x.

**Figure 8.**
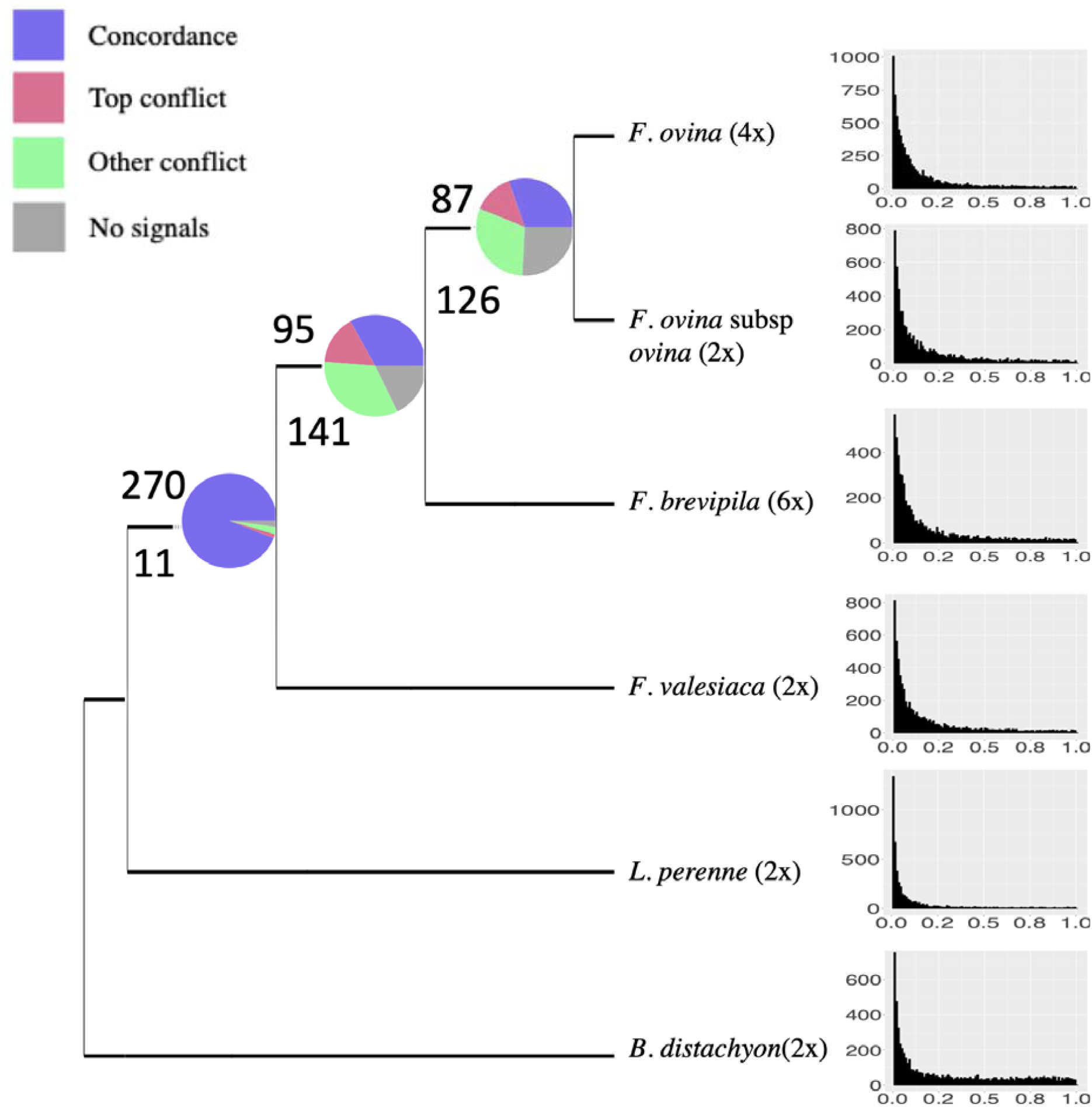
Species tree of *F. ovina* complex reconstructed by ASTRAL using 286 single-copy genes. Pie charts at nodes indicate the proportion of informative gene trees (bootstrap support of 50 or higher) that were concordant (blue) or conflict (top conflict in red and all other conflicts in green) with the species tree topology. Gray represents gene trees with low (< 50) bootstrap support. Total number of concordance gene trees were above the branch and the total number of conflicting gene trees were below the branch. Ks plots to the right of each tip showed no distinguishable Ks peak, suggesting the whole genome duplication events happened very recently.

## Discussion

In this study, we used the PacBio isoform sequencing platform to generate a reference transcriptome of *F. brevipila* using root, leaf, crown, and inflorescence tissue types. In the absence of a reference genome, we used multiple protein and pathway databases to annotate the reference transcriptome. We used Illumina sequencing data from *F. brevipila* leaf tissue to polish the PacBio reads and to generate a *de novo* assembly to evaluate the coverage of the reference transcriptome. Finally, we carried out phylotranscriptomic analyses to investigate the phylogenetic relationships among *F. brevipila* and its close relatives.

The analysis of the perennial ryegrass draft genome, a close relative of *F. brevipila*. identified 28,455 genes (Byrne et al., 2015). Since *F. brevipila* is hexaploid, we would expect a higher total number of genes; however, we only identified 38,556 non-redundant transcripts in our reference transcriptome, which could be the result of multiple factors. First, tissues were sampled while growing in greenhouse conditions, and we only sampled at one time point. Therefore, it is likely that we are missing a number of genes that are not expressed at that time point. Secondly, we did not perform library size selection, which could lead to the over presence of some highly abundant transcripts and fail to capture less frequent transcripts. Finally, homeologs may be difficult to distinguish from sequencing errors and heterozygosity, given the highly similar subgenomes of *F. brevipila* as suggested by the lack of any clearly distinguishable Ks peak in our Ks analysis (**Figure S1**). Future research should sequence more tissue types, stress conditions, and time points to provide a more comprehensive transcriptome.

Next, we used the Illumina short reads sequencing to evaluate the completeness of the *F. brevipila* reference transcriptome. Comparing our reference transcriptome based on PacBio Isoseq alone, Illumina *de novo*, and the merged assembly, the PacBio Isoseq transcriptome had the best N50 of 2,584 bp while *de novo* assembly only had N50 of 1,157 bp. The merged contigs had the highest BUSCO completes (89.2%), reduced fragmented BUSCO (6.8%), and the lowest missing BUSCO (4%). When mapping Illumina sequencing reads to the three assemblies, the combined assembly has the least unmapped reads, followed by PacBio Isoseq, and *de novo* assembly. The PacBio assembly has the best mapping quality (42.9% MAPQ >=30), followed by the *de novo*, and then the combined assembly. The combined assembly had the highest number of reads with MAPQ <3 (mapping to multiple regions of the transcriptome), which suggests that when combine multiple assemblies using cd-hit-est, the sequence identity threshold played a crucial role. In our case, we used a 95% sequencing identity cut off. By using a lower cut off, we would likely increase the unique mapping (MAPQ>=30); however, we would potentially lose homolog or paralog information due to the hexaploid nature of the taxon. Similarly, if we used a more stringent cutoff, we would likely maintain more sequence information but lose a greater number of unique reads. This is an inherent problem in polyploid transcriptomics, especially when using short reads. Future whole genome sequencing will help researchers to overcome these limitations. Therefore, we present the polished Isoseq dataset as our reference transcriptome, instead of using the hybrid assembly.

Although PacBio Isoseq has proven to improve the transcriptome quality, it has several limitations. First, it has a higher cost and lower sequencing coverage compared to the Illumina dataset. When comparing the BUSCO assessment result, the PacBio Isoseq dataset had 28.8% BUSCO genes missing while the short read *de novo* assembly had only 5.4% despite sequencing from a single tissue. In addition, at present PacBio sequencing has a higher error rate compared to Illumina platforms. However, these limitations could be reduced by increasing the number of SMRT cell for sequencing, improvement of sequencing chemistry, enzyme (Hi-Fi), and a combination of using Illumina short reads sequencing and long reads (Rhoads and Au, 2015; Wenger et al., 2019). Finally, despite being capable of recovering full-length transcripts, 83 out of the 988 BUSCO genes in our PacBio Isoseq dataset remain fragmented.

In addition to transcriptome annotation using a multiple protein database, we identified potential microRNA precursors (miRNA). When searching miRNA precursor transcripts in our dataset, we found MIR6179, MIR1119, and MIR2927 had the highest presence in the four tissue types sampled. Previous studies have shown that MIR1119 is essential in regulating plant drought tolerance in wheat and dehydration stress in barley (Kantar et al., 2010; SHI et al., 2018). MIR2927 has been shown to regulate both abiotic (salt) and biotic (virus) stress response in rice (Sanan-Mishra et al., 2009). The function of MIR6179 was found to associate with gametocidal action in wheat (Wang et al., 2018). It is possible that the high abundance of these miRNAs reflects the importance of the pathways being targeted in *F. brevipila*.

Finally, to test the phylogenetic affinity of *F. brevipila*, we used the transcriptome data to construct the phylogenetic relationship among closely related taxa in the *Festuca – Lolium* complex. Since the error rate of the polished PacBio Isoseq is approximately 1%, it is challenging to distinguish whether sequence differences at or below this percentage are due to read errors, heterozygosity, or different homeologs/paralogs. For this reason, we collapsed redundant transcripts using a minimal sequence identity of 99% when constructing the reference transcriptome. This step could lead to overestimation of the true number of single copy genes. However, it is unlikely to significantly affect the phylogenetic affinity of *F. brevipila* in downstream phylotranscriptomic analyses.

When comparing the phylogeny from maternal (chloroplast) vs. bi-parentally inherited (nuclear) genes, we noticed a conflict between the placement of *F. brevipila* (6x) and *F. valesiaca* (2x; **Figure 7**). Among nuclear gene trees (**Figure 8**), approximately 40% of informative gene trees supported the nuclear species tree topology, with no single dominant alternative topology. Distribution of synonymous substitutions per site (Ks) lacks any clear peak in any of our ingroup species (**Figure 8**), despite the detection of a known Ks peak around 0.8 in the outgroup *Brachypodium* (**Figure S1**) (Wang et al., 2015). This suggests that the polyploidy event happened relatively recently and may not be distinguishable from sequencing and assembly errors. Given the short internal branch length among *F. brevipila* (6x), *F. valesiaca* and *F. ovina* (2x and 4x), hybridization, incomplete lineage sorting, and sequencing and assembly artifacts may contribute to the conflict among the nuclear genes. Ideally, we could use ortholog and homologous group information to further investigate this event; however, our data is unable to distinguish among these potential sources of gene tree conflict due to the difficulty of phasing homeologs. Future studies will be focused on whole genome sequencing and screening additional taxa that are closely related to *F. brevipila* to identify affinities of subgenomes of this important hexaploid turfgrass taxon.

## Conclusion

In this study, we developed the reference transcriptome of *F. brevipila* using PacBio isoform sequencing and polished the reads and evaluated its completeness using Illumina sequencing. Distribution of synonymous distances among paralogs within *F. brevipila* suggested highly similar subgenomes that are difficult to distinguish from sequencing errors. In addition, we carried out phylotranscriptomic analysis to confirm the close phylogenetic affinity of *F. brevipila* with *F. ovina* and *F. valesiaca*, with high levels of gene tree discordance among them. This dataset provided insight into a complicated hexaploid transcriptome and laid the foundation for future genomics research on this important turfgrass taxon.

## Methods

### Plant Materials

*Festuca brevipila* genotype SPHD15-3 from the University of Minnesota breeding program (material used to develop the advanced breeding population MNHD-15) was used for this study (plant materials available upon request). A clonal population was vegetatively propagated and grown using BRK Promix soil (Premier Tech, USA) in the Plant Growth Facility at the University of Minnesota, St. Paul campus under 14 hour daylight and 8 hour darkness with bi-daily irrigation and weekly fertilization (906 grams of ammonium sulfate, 950 grams of Peat-lite 20-10-20 and 38 grams of Sprint 330 mixed into 5 gallons of water with a Dositron fertilizer injector set at a 1:100 ratio).

To capture a broad representation of the *F. brevipila* transcriptome using PacBio Isoform sequencing, we collected root, crown, leaf, and inflorescence tissues. Plants were vernalized in a 4L cold room for 3 months with 8 hour daylight and 16 hour darkness. Plants were then moved back to greenhouse conditions as described above with daytime temperature of 25L to recover for one month and allow inflorescence development. The root, crown, leaf, and inflorescence tissues were harvested from a two-month old plant and flash frozen for RNA extraction (**Figure 1**).

The species identity of plant materials used in this study were confirmed following guidelines described by Wilkinson and Stace (1991) using vein number, leaf morphology, and flow cytometry (Qiu et al., 2019). Voucher specimen for *F. brevipila* is deposited in the University of Minnesota Herbarium (Qiu 1, MIN). For the phylotranscriptomic analysis, *F. ovina* subsp. *ovina* (USDA PI 676177), *F. valesiaca* (PI 422463), *F. ovina* cv. Quatro (obtained from National Turfgrass Evaluation Program, http://www.ntep.org), and *F. brevipila* SPHD15-3 were grown in the greenhouse with the same condition described above. Leaf samples (2-3 cm above the leaf base) were harvested and flash frozen for RNA extraction.

### RNA Extraction and Sequencing

To maintain RNA integrity, RNA extractions for Isoseq were done using a Quick-RNA Miniprep Kit (ZYMO Research, Catalog number R1055) in a cold room following the manufacturer’s instructions. RNA extraction for Illumina sequencing was performed at room temperature using the same method. A Qubit Fluorometer (ThermoFisher Scientific) was used for initial RNA quantification, before an Agilent 2100 Bioanalyzer (Agilent) was used to assess the RNA integrity. Only samples with a RIN >8.0 were used for sequencing library construction.

For PacBio Isoform sequencing, four sequencing libraries (one for each tissue type) were constructed using the SMARTer PCR cDNA Synthesis Kit (ClonTech, Takara Bio Inc., Shiga, Japan) by tissue type with no size selection by NovoGene, China. Sequencing was performed on a PacBio Sequel II (PacBio, CA) instrument. Illumina sequencing libraries were constructed using a TruSeq® Stranded mRNA Library Prep kit (Illumina) and sequenced on an Illumina HiSeq 4000 instrument in 150 bp paired-end sequencing mode. All sequencing data generated from this study have been deposited in NCBI SRA under BioProject PRJNA598357.

### Illumina Sequencing Data Processing and *de novo* Assembly

Illumina adaptor sequences were trimmed using Trimmomatic with the default settings (v 0.32) (Bolger et al., 2014). Quality trimming was performed using the seqtk tool with -q 0.01 (https://github.com/lh3/seqtk). Trinity (v 2.4.0) was used to assemble transcriptomes of *F. ovina* subsp. *ovina, F. valesiaca, F. ovina* cv. Quatro, and *F. brevipila* (Grabherr et al., 2011) with max memory 62G, 24 CPUs, bflyCalculateCPU, and minimum contig size of 200 bp.

### PacBio isoform sequencing (Isoseq) Data Processing

The software SMRTlink (v 2.3.0, PacBio) was used to filter and process original sequencing files, with minLength 0, minReadScore 0.8 to produce the subreads file. To identify full-length transcripts, the Isoseq 3 (v 3.1.0, PacificBiosciences) pipeline was installed locally using conda (Anaconda-2.3.0) under bioconda following instruction by PacBio (https://github.com/PacificBiosciences/Isoseq_SA3nUP). Circular consensus sequences (ccs) bam files were generated using ccs command with --noPolish, --maxPoaCoverage 10, and 1 minimum passes options. To classify full-length (FL) reads, we identified and trimmed the 5’ and 3’ cDNA primers (primer sequences can be found in **Table S1)** in the ccs using lima with -- dump-clips, --no-pbi, --peak-guess option. Poly(A) tails in FL reads that were at least 20 bp long were removed with --require-polya. Bam files produced from the previous steps for four tissue types were merged into one dataset, and the source files were also merged into a subreadset.xml file. Clustered reads were polished using the Isoseq3 polish function to produce high-quality FL transcripts (expected accuracy ≥ 99% or QV ≥ 30).

To correct the PacBio Isoseq FL transcripts to obtain the reference transcriptome, high quality Illumina paired-end reads from *F. brevipila* leaf tissue were used in the LoRDEC -correct function in LoRDEC program (v 0.6) with configuration *k-mer* size = 19, solid-*k-mer* = 3, error rate 0.4, and maximum branch time 200 (Salmela and Rivals, 2014).

### Removing Sequence Contamination and Collapsing Redundant Sequences

To remove microbial sequence contamination in both Isoseq and Illumina *de novo* transcriptomes, bacterial and virus genomes were downloaded from NCBI ftp://ftp.ncbi.nih.gov/genomes/archive/old_refseq/Bacteria/all.fna.tar.gz, ftp://ftp.ncbi.nih.gov/genomes/Viruses/all.fna.tar.gz, a mapping index was built using bwa64 (v 0.5.9-r16) included in the deconseq program (v 0.4.3) (Schmieder and Edwards, 2011). Contaminant transcripts were removed using the deconseq.pl script in the deconseq program before performing downstream analysis.

Because the PacBio Isoseq3 pipeline output sequences with 99% accuracy, we used ‘cd-hit-est’ in the CD-HIT (v 4.8.1) package to cluster sequences based on sequence identity of at least 99% and the shorter sequence covered at least 99% of the longer sequence to collapse redundant transcripts in each transcriptome (-c 0.99 -G 0 -aL 0.00 -aS 0.99 -AS 30 -M 0 -d 0 -p 1 [Isoseq transcriptome]; -c 0.98 -p 1 -T 10 [*de novo* assembly]) (Li and Godzik, 2006).

### Transcriptome Completeness Analysis

To evaluate the completeness of the assembled *F. brevipila* reference transcriptomes, we used 1335 core embryophyte genes (embryophyta_odb10) from Benchmarking Universal Single-Copy Orthologs (BUSCO v3) (Simão et al., 2015). Besides evaluating the completeness of the Isoseq reference transcriptome, we also evaluated the completeness of the *de novo* transcriptome separately, and the combined transcriptome (merged Isoseq transcriptome and short read *de novo* transcriptome using cd-hit-est to collapse redundant sequence based on 95% sequence identity) (Li and Godzik, 2006).To evaluate how the combined assembly could improve short read mapping, we mapped RNA sequencing reads used for *de novo* assembly to the individual and the combined assembly using bowtie2 program (Langmead and Salzberg, 2012). Mapping statistics were summarized using the samstat tool (v 1.5.1) (Lassmann et al., 2010).

### Transcriptome Functional Annotation

The reference transcriptome was searched against NCBI NR protein and SwisProt, UniProt protein database using diamond BLASTx (v 0.9.13) with e-value < 1e-5. Kyoto Encyclopedia of Genes and Genomes (KEGG) pathway analyses were performed using the KEGG Automatic Annotation Server (KASS, https://www.genome.jp/kegg/kaas/). *Oryza sativa, Brassica napus, Zea mays*, and *Arabidopsis thaliana* were set as references with a single-directional best hit model. Gene Ontology (GO) terms were produced by the interproscan program (Quevillon et al., 2005).

### Identification of lncRNAs and *miRNA* from PacBio Sequences

The prediction of long non-coding RNAs (lncRNAs) was performed using an improved *k-mer* scheme (PLEK) tool (v 1.2) with a minimum sequence length of 200 bp (Li et al., 2014). Putative miRNA precursors were identified by BLASTn of reference transcriptome sequences against the plant miRNAs database (http://www.mirbase.org) that includes both hairpin and mature miRNA with e-value δ 10^−5^ (Kozomara and Griffiths-Jones, 2013). For each sequence the BLASTn hit with the highest bit score was kept.

### Phylotranscriptomic Analyses of *Festuca-Lolium* Taxa

To investigate the nuclear phylogeny of *F. brevipila* and close related species, we also generated the transcriptome for three additional taxa (*F. ovina* subsp. *ovina, F. valesiaca*, and *F. ovina*) in this study using *de novo* assembly method described above. In addition, we included four publicly available transcriptomes from closely related species selected according to previous phylogenetic analyses based on chloroplast sequences (**Table 1**) (Hand et al., 2013; Qiu et al., 2019). Protein and coding sequences of all *Festuca – Lolium* taxa were predicted using the TransDecoder program (https://github.com/TransDecoder) (Wu and Watanabe, 2005). Proteome from genome annotation of *Brachypodium distachyon* were included as the outgroup.

Orthology inference was carried out using protein sequences of all nine taxa in OrthoFinder 2 (Emms and Kelly, 2019). Single-copy protein sequences were aligned using the MUSCLE program (v 3.8.31) with the parameters -maxiters 16 -diags -sv (Edgar, 2004). To improve the phylogenetic resolution among closely related taxa, we used the PAL2NAL program (v 14) to align coding sequences according to the protein alignment (Suyama et al., 2006). The resulting coding sequence alignments were trimmed using TrimAl with the parameters -gt 0.9 - cons 60, which removes all positions in the alignment with gaps in 10% or more of the sequences and prints the 60% best positions (with fewer gaps) (Capella-Gutiérrez et al., 2009).

To trace the maternal relationships of the focal taxa, chloroplast gene sequences for *F. ovina* subsp. *ovina, F. valesiaca*, and *F. brevipila* were extracted from *de novo* assembled transcriptomes by aligning assembled contigs to the *F. ovina* reference chloroplast genome (MN309824) using Burrows-Wheeler Aligner (Li and Durbin, 2009). Sequences for an additional five closely-related taxa were obtained from NCBI by BLASTn using the *F. ovina* chloroplast genome sequences as the query. The *Brachypodium* chloroplast sequence was used as the outgroup (**Table 1**). Three chloroplast intergenic spacer regions that covered *rps2-atpA, rpl32*-*ccsA*, and *clpP*-*psbH* region had the highest completeness and taxon representation and were each aligned using MAFFT [v 7; (Katoh and Standley, 2013)] before concatenated into the chloroplast supermatrix.

Phylogenetic trees were constructed using the RAxML program (v 8.2.12) for both the concatenated nuclear and chloroplast supermatrices under the GTR+GAMMA model with 1,000 bootstraps (Stamatakis, 2006). The phylogenetic trees were visualized using FigTree v 1.4.3 (Rambaut, 2012). Because the chloroplast sequences and transcriptomes for *L. perenne, L. multiflorum, F. pratensis, F. arundinacea, B. distachyon* were downloaded from NCBI/EBI and they might not come from the same genotypes, for these taxa, we present the nuclear and chloroplast phylogenies separately without carrying out conflict analyses.

Next, we further explored phylogenetic conflict among chloroplast and nuclear gene trees conflicts among *F. ovina* subsp. *ovina, F. ovina*, and *F. valesiaca* using transcriptomes generated in this study and the Isoseq transcriptome for *F. brevipila. Lolium perenne* and *B. distachyon* were included as outgroups. We carried out a new OrthoFinder run with a reduced set of taxa and used single-copy genes predicted from the new run. The coding sequence alignment for each single-copy gene was generated using the methods described above. Gene trees were estimated using the RAxML program (v 8.2.12) under GTR+GAMMA model with 1000 bootstrap. A species tree was estimated using ASTRAL v 5.6.3 using gene trees from the previous step (Mirarab et al., 2014). The concordance and conflicts among gene trees were visualized using Phypart (Smith et al., 2015) by mapping single-copy gene trees rooted by *Brachypodium distachyon* to the rooted species tree from ASTRAL. Tree concordance and conflicts were plotted using the python script phypartspiecharts.py available at https://github.com/mossmatters/phyloscripts/tree/ master/phypartspiecharts (last accessed on Feb 2, 2020).

For the 6 species in the ASTRAL tree, the distributions of synonymous substitutions per site (Ks) within a genome was calculated using wgd program (v 1.1.1, https://github.com/arzwa/wgd) (Zwaenepoel et al., 2019). Ks value was visualized in R (R core, 2013).

## Declarations Abbreviations

Full-length transcripts: FL transcript
PacBio Isoseq: PacBio Isoform Sequencing
NR-FL: Non-redundant full length
BUSCO: Benchmarking Universal Single-Copy Orthologs
KEGG: Kyoto Encyclopedia of Genes and Genomes
GO: Gene Ontology
lncRNAs: Long non-coding RNAs
ASTRAL: Accurate Species Tree Algorithm
NR protein: Non-Redundant Protein

## Declarations

### Ethics Approval and Consent to Participate

Not applicable

### Consent for Publication

Not applicable

### Availability of Data and Material

All PacBio IsoSeq and Illumina RNA-Seq data generated for this work were deposited at NCBI under the BioProject PRJNA598357. The raw PacBio reads of inserts are available with the SRA accession numbers SRX7658820–SRX7658823. Illumina RNA sequencing reads are available with the SRA accession numbers SRX7658307–SRX7658309, and SRX7656997. The PacBio transcriptome and annotation had been deposited at https://conservancy.umn.edu/handle/11299/211413 for the community access. Plant materials are available from the UMN turfgrass breeding program.

## Authors’ Contributions

Y. Q. performed the experiments, analyzed the data, and wrote the manuscript; Y. Y. helped with phylogenetic analysis; C. D. H. helped with the genomics analysis; E. W. secured funding for this project, supervised this research. All authors contributed to the revising of the manuscript and approved the final version.

## Acknowledgments

The authors would like to thank Minnesota Supercomputing Institute for providing computational resources.

## Figures and Tables

**Figure S1.**
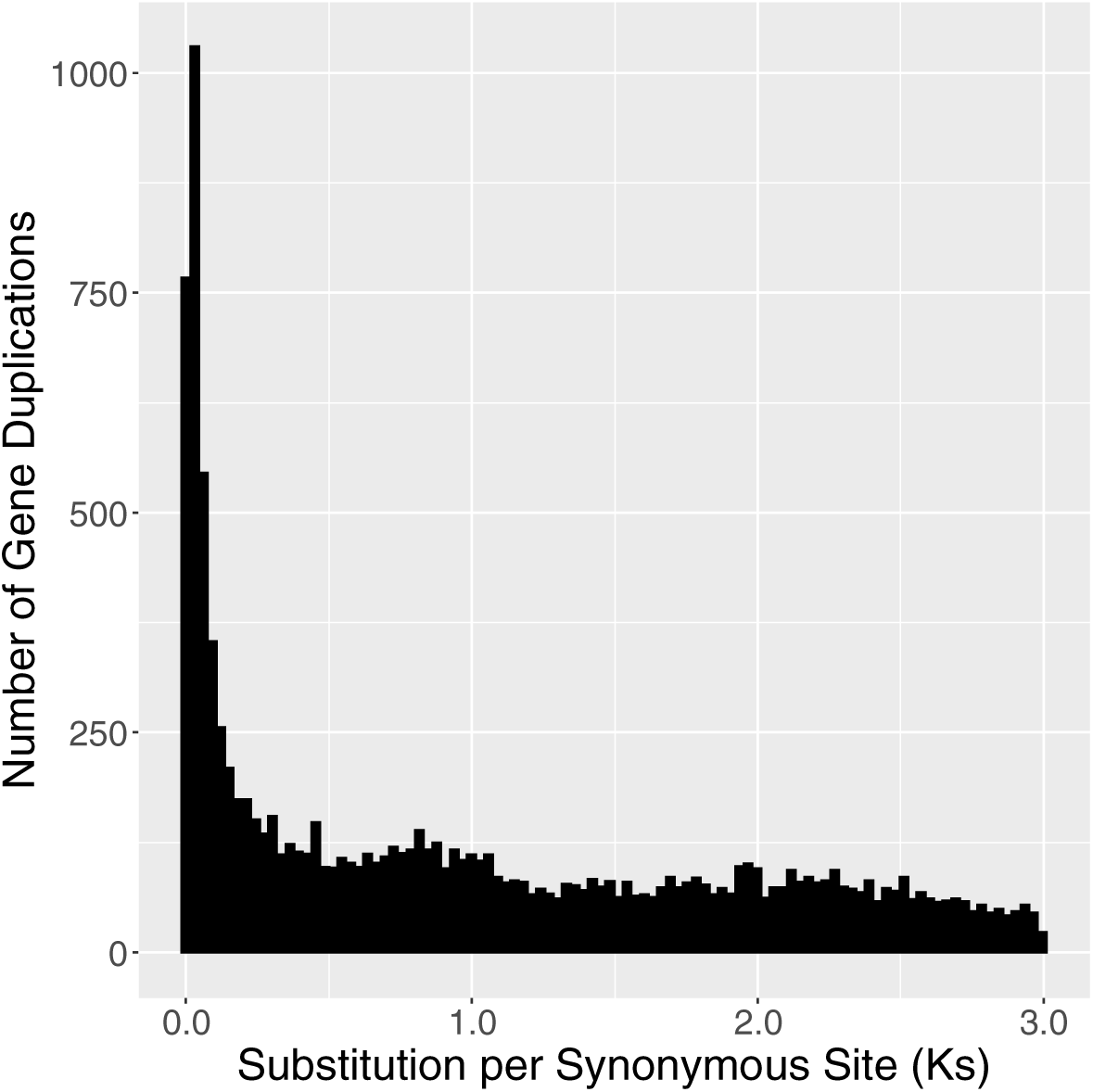
Ks analysis of the outgroup *Brachypodium distachyon* proteome from genome annotation, showing the known Ks peak at around 0.8.

**Table S1.**
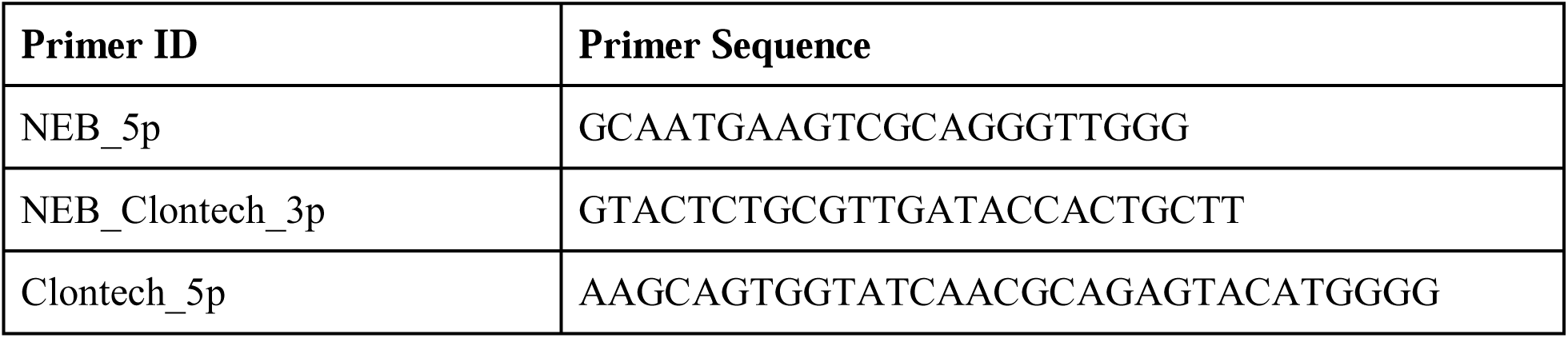
Primer sequences used for ccs file trimming using the Lima program

**Table S2.**
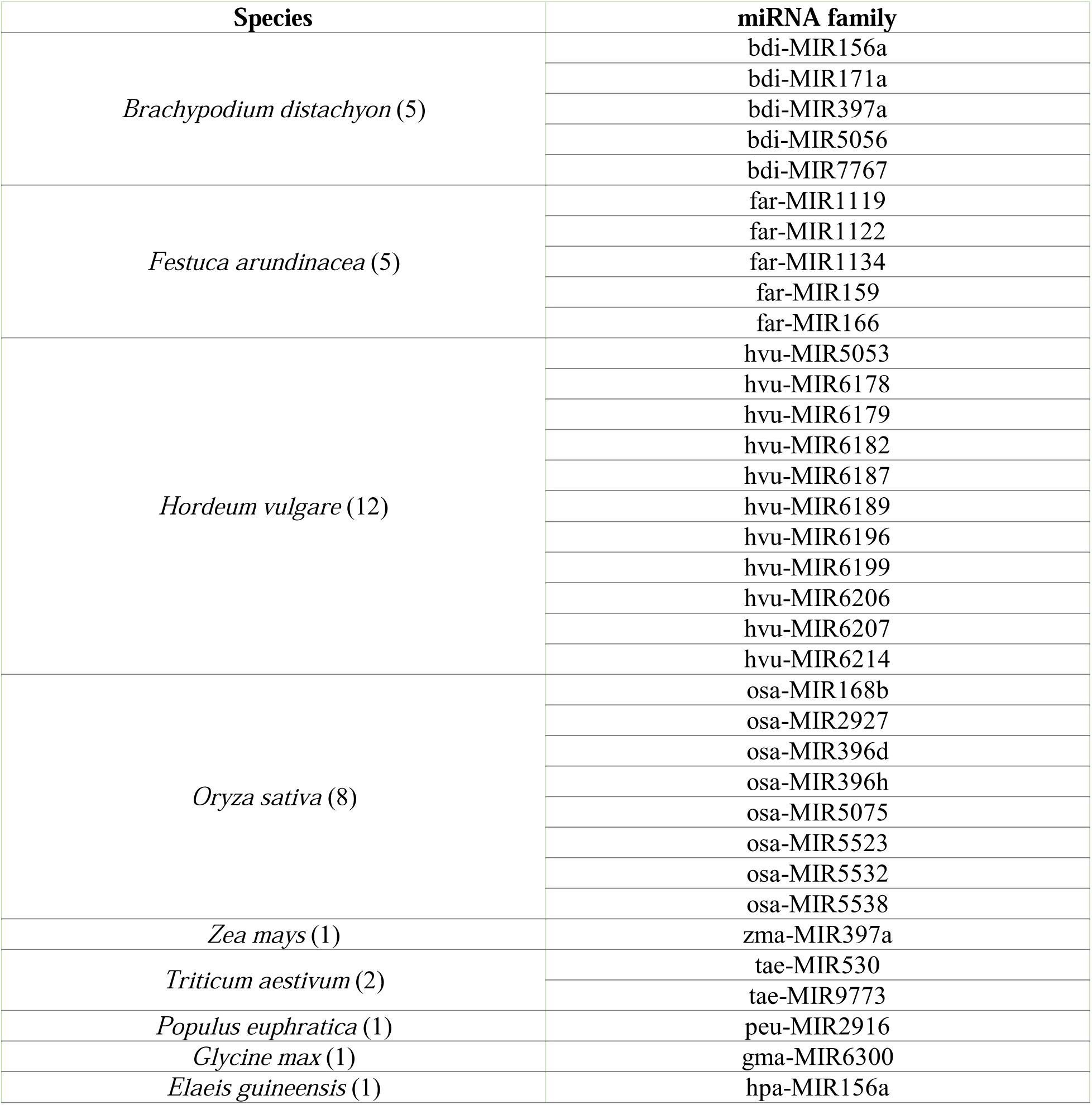
miRNA gene families identified in *F. brevipila* NR transcriptome.

